# 16S rRNA Survey Reveals the Potential of Oral Microbiota in Distinguishing Patients with Chronic Heart Failure from Healthy Controls

**DOI:** 10.1101/2025.08.12.669863

**Authors:** Xing Ye, Jie Jia, Xiaotao Jiang, Jiao Jiao Li, Yanlin Zeng, Yan Wang, Zhongqi Yang, Tianhui Yuan, Jing Sun

**Author notes:** Corresponding authors. Tianhui Yuan: The First Affiliated Hospital of Guangzhou University of Chinese Medicine, No.12, Ji Chang Road, Baiyun District, Guangzhou, 510405, China. E-mail addresses. (T. Yuan).

## Abstract

**Background:** Oral microbiota can reflect physiological functions and pathological conditions in human body. Patients with chronic heart failure (CHF) exhibit distinct oral health status compared to healthy controls (HCs), which is attributed to the differences in dominant microbial communities present in the oral cavity. Up to date, there are few studies examined the association between CHF and dominant oral microbiota. To fill in this research gap, this study aimed to investigate the differences of oral microbiota between CHF patients and HCs, to identify valuable novel biomarkers for CHF.

**Methods:** Chronic heart failure patients and healthy volunteers were recruited. Oral microbiota samples were then collected using oral swabs, and 16S rRNA sequencing was employed to analyze the microbiota. Statistical analysis was conducted to identify key bacteria at multiple taxonomic levels in the oral microbiota samples from both the CHF patient and healthy control groups, with a focus on core genera to identify potential biomarkers and evaluate their diagnostic efficacy.

**Results:** There were 60 CHF patients and 30 HCs were recruited, with 42 CHF patients with New York Heart Association (NYHA) functional class II-IV and 28 HCs were included in the final analysis. The alpha diversity was higher in HCs, while beta diversity was higher in CHF patients. The CHF patients showed significant differences from HCs at five gene (phylum, class, order, family and genus) levels by analyzing the relative richness of microbiota at different taxomal levels. Altogether 14 microbes could distinguish CHF patients from HCs, i.e., *Abiotrophia*, *Butyrivibrio*, *Lactobacillus*, *Capnocytophaga* and *Neisseria* which are more abundant in CHF patients, and *Actinomyces*, *Anaerovorax*, *Eubacterium*, *Kingella*, *Mogibacterium*, *Peptococcus*, *Peptostreptococcus*, *Solobacterium* and *TM7_genus_incertae_sedis* which are more abundant in HCs. Furthermore, the AUC of their combined diagnosis was 83.7% (95% confidential interval 74.1%–93.3%), which have high reliability for the diagnostic significance. In accordance to Spearman’s correlation, *Eubacterium, Solobacterium* and *Rhizobium* were core genera and the abundance of *Eubacterium* and *Solobacterium* exhibited downward trends as NYHA class increases.

**Conclusion:** This study revealed the dysbiosis of the oral microbiota in CHF patients and identified potential biomarkers for CHF diagnosis and management.

**Graphical abstract:** 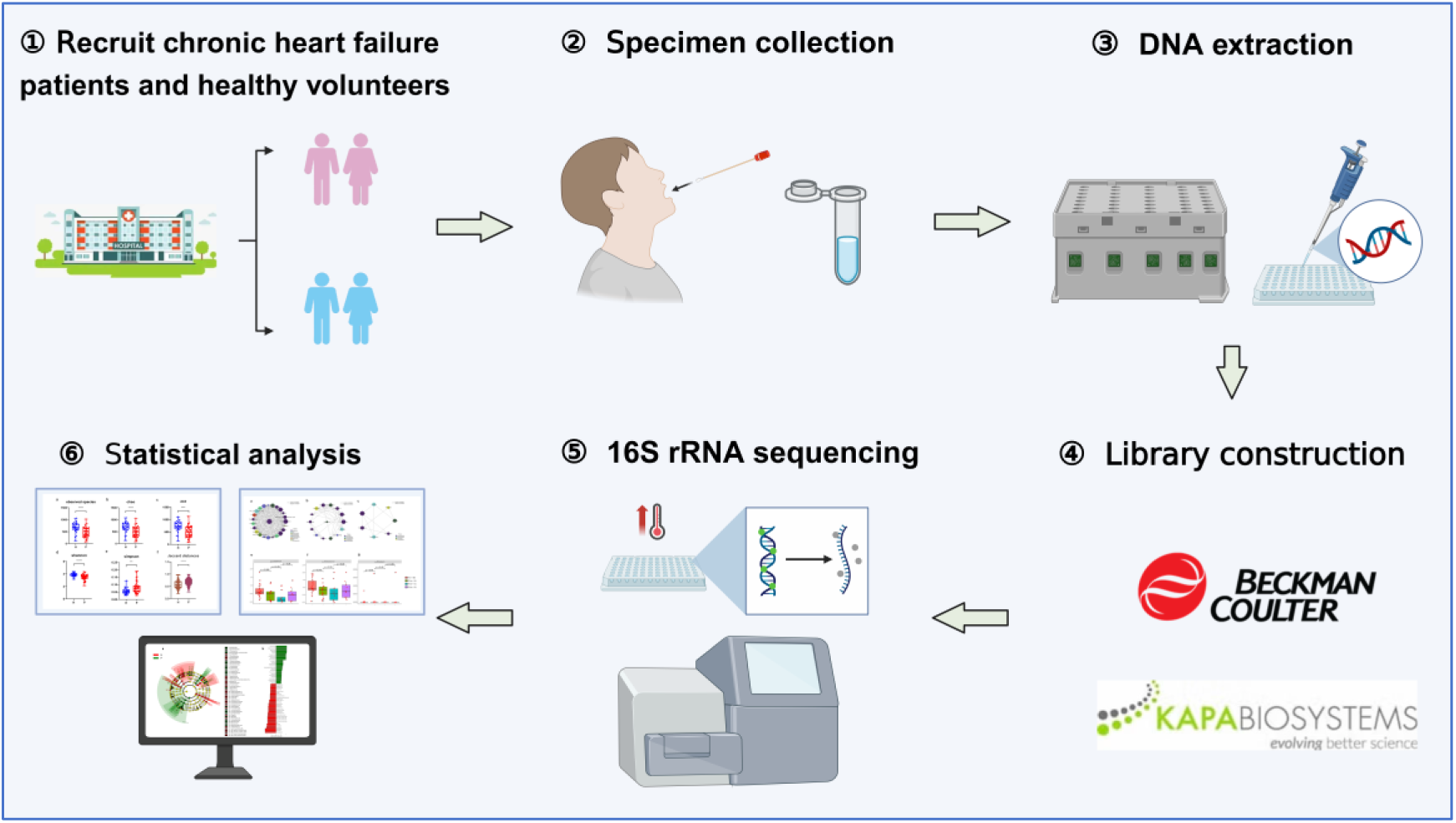

## 1. Introduction

Chronic heart failure (CHF) is one of globally major public health problems[1]. As of 2017, an estimated 64.3 million people worldwide suffering from heart failure, with an estimated 4.2 million patients in China[2, 3]. The consequences of CHF are severe, often leading to reduced quality of life, frequent hospitalizations, and high mortality rates. The prevalence of heart failure and costs of managements are expected to increase with the population aging and rising incidence of cardiovascular diseases[4, 5]. Despite advancements in treatment, accurate and timely diagnosis remains a challenge due to the lack of effective and specific biomarkers. Therefore, utilization of biomarkers in diagnosis and prognosis of heart failure is increasingly recognised[6, 7]. However, the current evidence in the biomarkers associated with heart failure is mixed and insufficient[8]. Currently available diagnostic biomarkers, such as B-type natriuretic peptide (BNP) and N-terminal pro-BNP (NT-proBNP), though useful, often lack specificity and may require invasive operations for comprehensive assessment. Their limitations in predicting major clinical events indicate a continuous need for more effective biomarkers[9, 10].

Previous studies have reported close relationships among the human microbial community, human physiology and pathology[11, 12]. For instance, intestinal microbiota, as a prominent area of scientific investigation, has been implicated in the pathogenesis of numerous disease entities. In a study conducted by Hu et al, intestinal flora is expected to become a new target for nerve protection through many pathways in post-stroke injury rehabilitation[13]. Wang’s study demonstrated that the intestinal microbiota plays a crucial role in modulating the maturation of the host immune system, functions as both a diagnostic biomarker and therapeutic response predictor in cancer immunotherapy, and significantly contributes to the pathogenesis of colorectal carcinoma[14]. In addition, previous studies have reported that the gut microbiota plays a significant role in the pathogenesis and progression of heart failure (HF), indicating its potential diagnostic and therapeutic value in the clinical management of cardiovascular disorder[15]. While much attention has been given to the gut microbiota, interestingly, many systemic conditions manifest in the mouth, highlighting the oral cavity’s role as an indicator of systemic disturbances, thus the oral microbiota offers unique advantages for biomarker discovery. The study from Georges et al.[16] revealed a strong correlation between dysbiosis in the oral environment and the pathogenesis of systemic diseases. And Reis et al. found that it potentially interacts with intestinal microbes and influencing systemic environments[17]. The oral microbial community occupies the most anterior position in the human digestive system, and the oral microbiota shares considerable features with the gut microbiota both in mucosal epithelia and microbial diversity[18, 19]. But unlike gut microbiota sampling, which often requires fecal or blood sample, oral microbiota can be easily and non-invasively collected, making it a practical choice for clinical applications. Moreover, it offers a personalized microbial signature for each patient. This convenience and diagnostic potential position the oral microbiota as an underexplored yet valuable method for biomarker research.

In recent years, oral microbiota has been increasingly utilized to aid in disease diagnosis and biomarker discovery. Studies have demonstrated its utility in identifying microbial alterations associated with aspiration pneumonia[20], pancreatic cancer[21], chronic kidney disease[22], oral cancers[23], chronic hepatitis[24], and gastrointestinal disorders[25]. In addition, a prospective multicenter study has proved it can be useful in determining the benign or malignant status of indeterminate pulmonary nodules (IPN)[26]. However, despite these advancements, the diagnostic potential of oral microbiota in CHF remains unexplored. This gap in research highlights an opportunity to investigate its role as a non-invasive diagnostic tool for CHF.

To investigate the composition of oral microbiota in CHF patients, we employed 16S rRNA gene sequencing, a widely used and highly reliable method for microbial community analysis. 16S rRNA sequencing has several distinct advantages, including exceptional sensitivity, remarkable specificity, and superior cost-effectiveness, enabling precise identification and comprehensive characterization of microbial communities, particularly in challenging low-biomass samples, widely used in the study of intestinal microbes[27, 28], which has been successfully applied in studies exploring microbiota in various diseases, such as inflammatory bowel disease[29] and type 2 diabetes[30], demonstrating its robustness and utility in clinical microbiome research.

This study was designed to conduct a comprehensive analysis of oral microbiota in chronic heart failure (CHF) patients and healthy controls (HCs) through 16S rRNA sequencing coupled with innovative bioinformatics approaches. The primary objectives were to elucidate the potential association between oral microbiota and CHF pathogenesis, identify distinctive microbial signatures specific to CHF, explore novel non-invasive biomarkers, and ultimately contribute to the development of innovative diagnostic strategies for CHF management. The detailed study design process is shown in graphical abstract.

## 2. Materials and Methods

### 2.1 Sample collection

Sixty patients with CHF in the First Affiliated Hospital of Guangzhou University of Chinese Medicine and Guangdong Provincial Hospital of Chinese Medicine were recruited from January 1st, 2018 to January 1st, 2021. Adults (18 years or older) with a confirmed diagnosis of CHF according to guidelines for the diagnosis and management of chronic heart failure were recruited[31]. Thirty healthy volunteers aged >18 without cardiovascular disease, kidney disease, tumor and other chronic diseases from Physical Examination Center of the two hospitals were matched to the CHF patients in age and gender. Patients or healthy volunteers were excluded if they (1) had oral, tongue or dental disease; (2) suffered from upper respiratory tract infection in the past four weeks; (3) have used antibiotics and immuno-suppressants in the past four weeks; (4) were pregnant or in lactation; (5) patients who were life-threatening, transferred from intensive care or have received chemotherapy and radiotherapy before admission would excluded. The investigation conform to the principles outlined in the Declaration of Helsinki. All participants provided written informed consent, and the Institutional Ethics Committee granted approval for the protocol (No. K 【2020】 061).

### 2.2 Specimen collection, DNA extraction and preservation

For each biological replicate, oral microbiota samples were collected from participants in the morning prior to tooth brushing and breakfast. The oral mucosa was scraped 5 to7 times by oral swabs and the coat samples were stored in a 1.5 ml Eppendorf tube with phosphate-buffered saline. After the collection, the tubes were centrifuged at 5000 rpm for 10 min. The supernatant was discarded whereas the sediment was preserved and stored at -80 °C before analyzing. Microbial DNA of the oral microbiota was extracted by QIAamp® DNA Mini and Blood Mini Handbook Kit (Qiagen, Hilden, Germany) according to instructions.

### 2.3 DNA quality inspection, Library construction and sequencing

DNA extracted should meet following requirements: (i) concentration is more than or equal to 40ng/μl; (ii) total quantity of DNA is more than or equal to 400ng; (iii) OD260/280 ranged from 1.8 to 2.0. The library was constructed via Beckman Coulter Genomics, KAPA Biosystems. We amplified the v3 and v4 region of 16SrDNA and added adapter as well as index using PCR. Magnetic bead was adopted to purify the products. After library passing quality control, libraries were sequenced on the Illumina MiSeq 2000 platform and the read length was set to 2X300bp.

### 2.4 Quality control and Assembly

We differentiated sequencing data for each sample according to the barcode sequence. Contaminated sequence of joints during sequencing, reads with low quality and certain content and reads containing N-base were removed. The paired reads are spliced using the make. contigs function in the mothur software package[32], and the paired reads obtained by double-end sequencing are assembled into a sequence using overlapping relationships. Therefore, Tags with hypervariable region were obtained. In order to retain better quality tags, the filter conditions were set as follows: (i) the minimum matching length was 10% of the length of the reads, i.e., 100 bp Paired End reads must have a minimum overlap length of 10 bp or more; (ii) The overlap area matching rate was 90%, i.e., the 10 bp overlap allows only one mismatch; (iii) N-base was not allowed to exist; (iv) The length was greater than or equal to 300 bp.

### 2.5 Operational taxonomic units (OTUs) and taxonomy profiling

After chimeric sequences were removed via USEARCH software (version 9.2)[33], operational taxonomic units (OTUs) were classified based on 97% similarity. Each high-quality sequence was compared against the Silva database to find the most similar specie information with more than 80% confidence. In order to obtain the taxonomic information of each OTU, all sequences in each OTU at 97% similarity level were analyzed by consistency. The species information of the OTU is determined according to the nearest ancestors of different sequences in the same OTU.

### 2.6 Statistical analysis

Alpha diversity was calculated using the following indexes: observed species (Obs), chao1 estimator, ACE estimator, Shannon indexes and Simpson indexes. Beta diversity was represented by Jaccard distance between samples. The comparison of alpha diversity index, beta diversity and microbes at different level were tested by Mann–Whitney U test. The correlation between species was analyzed using Spearman’s correlation and core genera were identified based on the maximum value of degree. LEfSe was applied to identify key bacteria in oral microbiota samples from CHF patient group and HC group at multiple levels and the results were visualized using taxonomic bar charts and cladograms. Receiver operating characteristics (ROC) analysis was adopyed to find biomarkers with diagnostic significance and evaluate their diagnostic efficacy. The difference was statistically significant when *P*<0.05.

## 3. Results

### 3.1 Sample of participants

There were 18 participants out of the 60 CHF patients were excluded since they have taken antibiotics within four weeks. And two participants in HC group were excluded as one was diagnosed with upper respiratory tract infection and the other one had poor DNA purity. The final sample consisted of 42 oral microbiota samples from CHF patients and 28 from HCs were obtained. Table 1 in appendices displayed the demographic characteristics of CHF patient group and HC group. Besides, distribution of NYHA class in CHF patients was shown in Fig.1, which were 19, 13, and 10 patients classified as NYHA class II, III, and IV, respectively.

**Fig.1.**
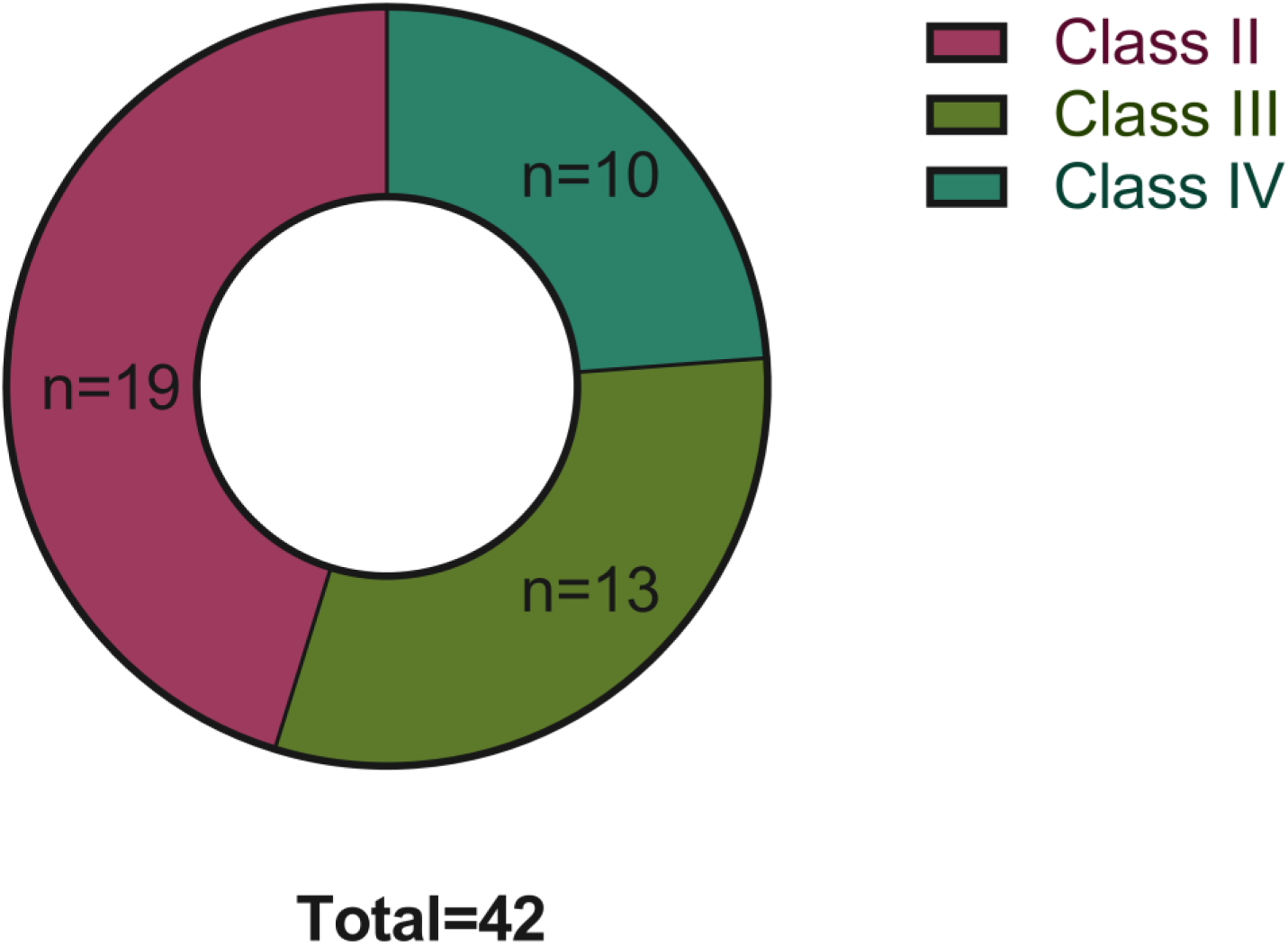
Distribution of NYHA class of CHF patients *n*=42.

**Table 1.** Demographic characteristics of CHF patients and healthy controls.

|  | Characteristics | Healthy controls | CHF patients |
| --- | --- | --- | --- |
| Sample size | / | 28 | 42 |
| Age | Mean ± SD | 26.25 ± 3.80 | 74.00 ± 13.62 |
| Sex | Male/Female | 12/16 | 18/24 |
| Disease duration | Mean ± SD | NA | 2.90 ± 2.01 |
| Hospital length of stay | Mean ± SD | NA | 5.43 ± 3.83 |

### 3.2 Differences of diversity of oral microbiota between CHF patients and HC

In order to comprehensively evaluate the overall diversity of microorganisms, we analyzed alpha diversity and beta diversity, which are commonly used in ecology to describe the diversity of species within and between habitats. In terms of alpha diversity, the abundance indexes including the Obs, the chao1 estimator and the ACE estimator, showed significant decreases in CHF patient group when compared with those of HC group (*P*<0.0001, *P*<0.0001, *P*<0.001), indicating that there is less species of microbiota in CHF patients (Fig.2a-c). Also, the Shannon indexes were significantly lower and the Simpson indexes were significantly higher in CHF patient group than those of HC group (*P*<0.0001, *P*<0.01), which revealed a more uniform species distribution in CHF patients (Fig.2d-e). As for beta diversity, the Jaccard distances between CHF patients were higher than those between HC(*P*<0.0001), reflecting that there are significant internal differences in oral microbiota between CHF patients and HC group (Fig.2f).

**Fig.2.**
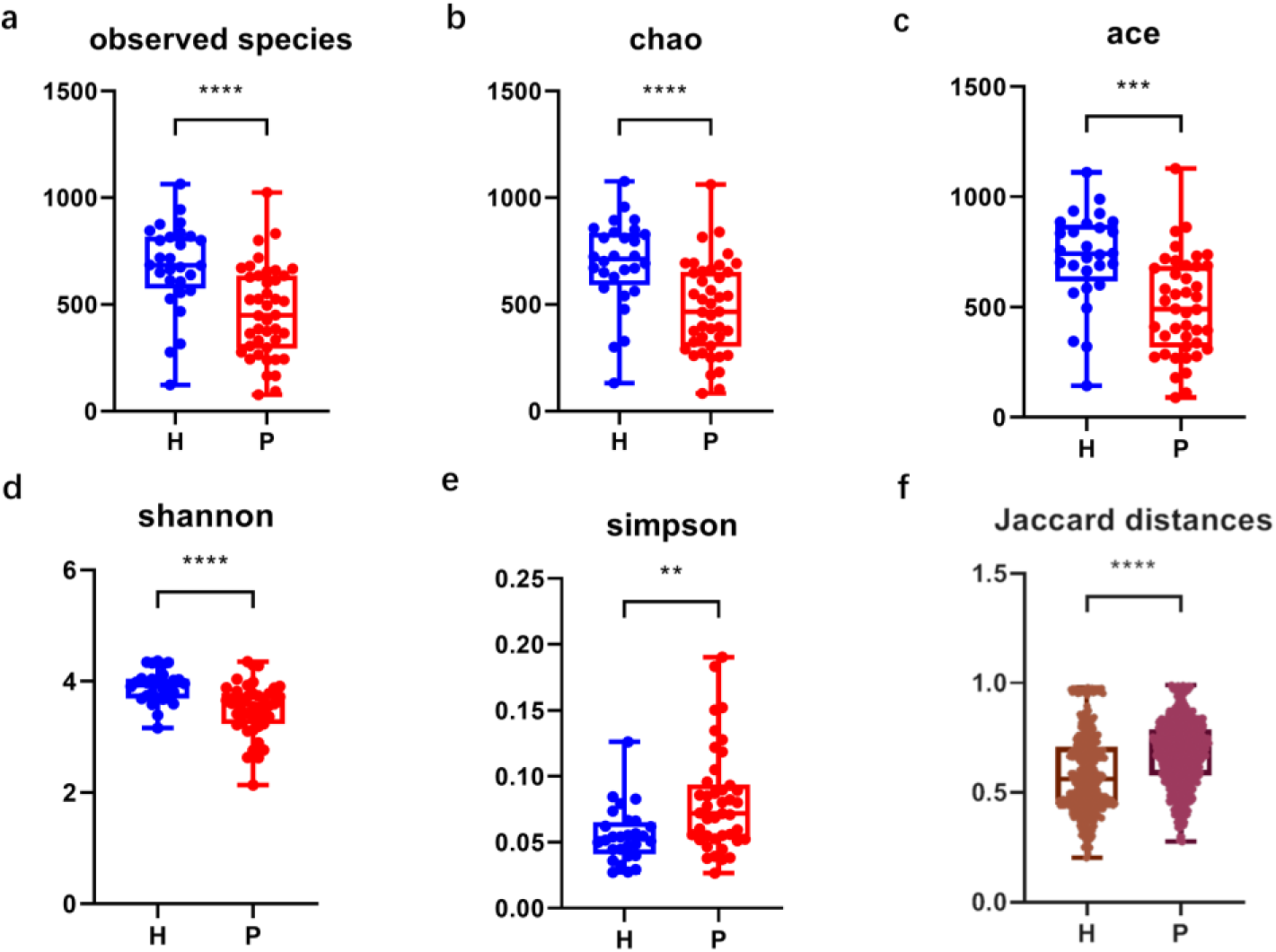
Diversity of tongue coating microbiota in CHF patients and HCs. **a**: Quantity of observed species of CHF patients and HCs; **b**: Chao indexes of CHF patients and HCs; **c**: Ace indexes of CHF patients and HCs; **d**: Shannon indexes of CHF patients and HCs; **e**: Simpson indexes of CHF patients and HCs; **f**: Jaccard distances of CHF patients and HCs. **Note**: H= HCs/heathy controls (*n*=28), P= CHF patients (*n*=42).

### 3.3 Different oral microbiota between CHF patients and HCs

To investigate the evenness and divergence of oral microbial community in different groups, we analyzed the relative abundance of microbiota at different taxomal levels and found that CHF patients showed significant differences from HCs at five gene (phylum, class, order, family and genus) levels. The main composition of the microbiota at phylum level was shown in Fig.3a. According to Mann-Whitney U test, the relative abundance of *Proteobacteria* were significantly higher in CHF patient group (*P*<0.05) (Fig.3b) while *TM7* was lower (*P*<0.05) (Fig.3c). At the class level (Fig. 3d), the relative abundance of *Erysipelotrichia, SR1_class_incertae_sedis, Gammaproteobacteria, TM7_class_incertae_sedis, Clostridia, Bacteroidia* and *Negativicutesin* the CHF patients were higher compared with HCs, whereas *Actinobacteria, Epsilonproteobacteria, Flavobacteria, Betaproteobacteria* and *Bacilli* were lower. At the order level, the relative abundance of *Bacteroidales, Clostridiales, Erysipelotrichales, Selenomonadales* and *TM7_order_incertae_sedis* in the CHF patients were higher, while A*ctinomycetales, Bacillales, Campylobacterales, Flavobacteriales* and *Lactobacillales* were lower (Fig. 3e). At the family level (Fig.3f, Fig.4), in CHF patient group, *Neisseriaceae, Flavobacteriaceae, Aerococcaceae, Bacteroidaceae, Bacteroidales_incertae_sedis, Comamonadaceae* and *Lactobacillaceae* were more abundant but the relative abundance of *Actinomycetaceae, Clostridiales_Incertae_Sedis_XIII, Erysipelotrichaceae, Eubacteriaceae, Peptococcaceae_1, Peptostreptococcaceae, Prevotellaceae* and *TM7_family_incertae_sedis* were lower. At the genus level (Fig.3g, Fig.5), there were 19 principally different genera of microbes. The microbes with higher abundance in CHF patient group included *Abiotrophia, Anaeroglobus, Bacteroides, Butyrivibrio, Capnocytophaga, Lactobacillus, Neisseria* and *Phocaeicola*. On the contrary, the other 11 microbes were less abundant in CHF patient group including *Anaerovorax, Eubacterium, Kingella, Mogibacterium, Murdochiella, Peptococcus, Peptostreptococcus, Simonsiella, Solobacterium* and *TM7_genus_incertae_sedis.* Moreover, LEfSe (Linear discriminant analysis effect size) were adopted to visualize the results (Fig.6a, 6b).

**Fig.3.**
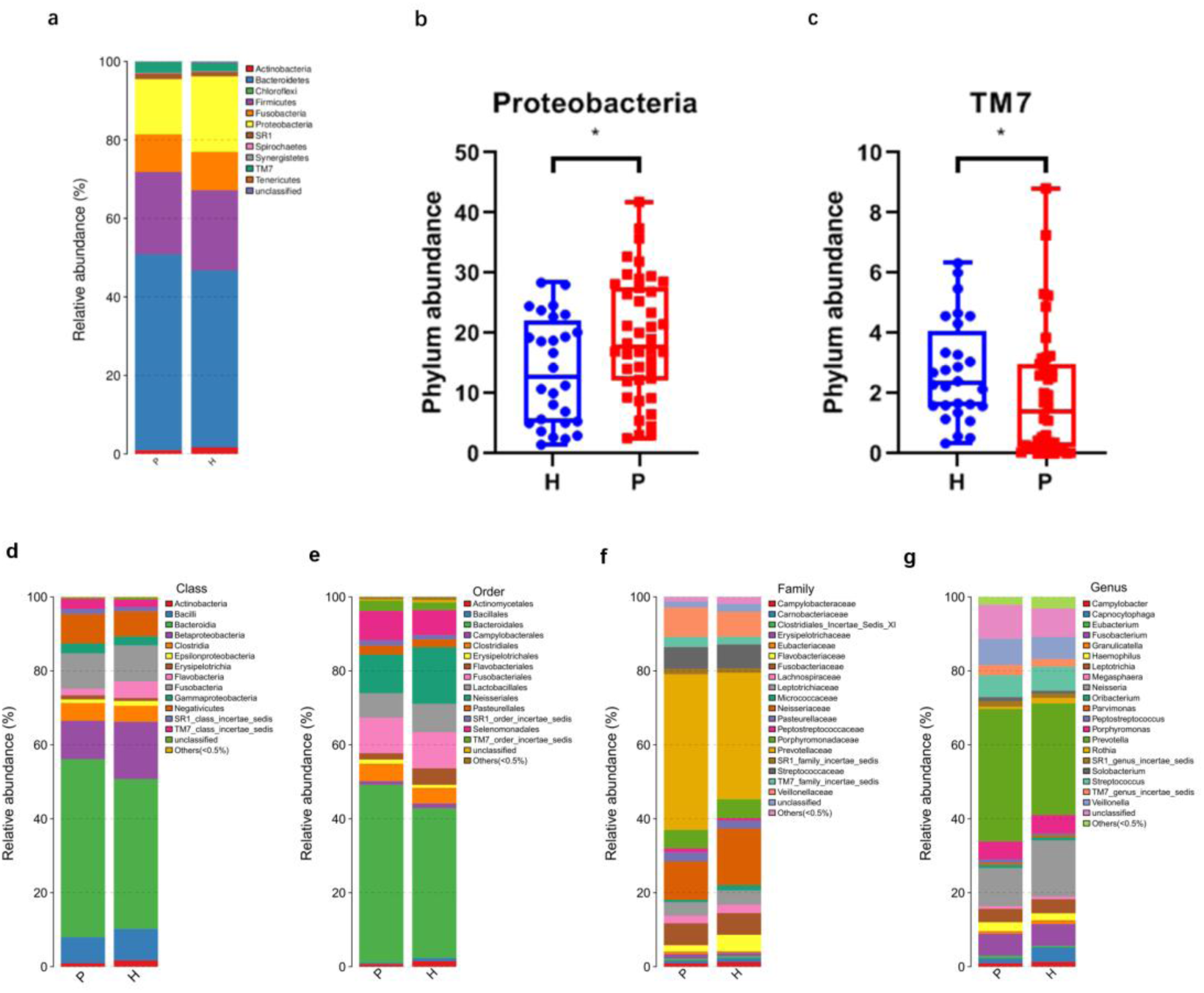
Composition of microbiota at phylum level (**a**) and microbes with significantly different abundance (**b,c**).The relative taxa abundance between HCs and CHF patients at class (**d**), order (**e**) family (**f**) and genus (**g**) level. **Note**: \**P* < 0.05; \*\**P* < 0.01; \*\*\**P* < 0.001 and \*\*\*\**P* < 0.0001; Comparison using the Mann–Whitney U test(b-g). H= HCs/heathy controls (*n*=28), P= CHF patients (*n*=42).

**Fig.4.**
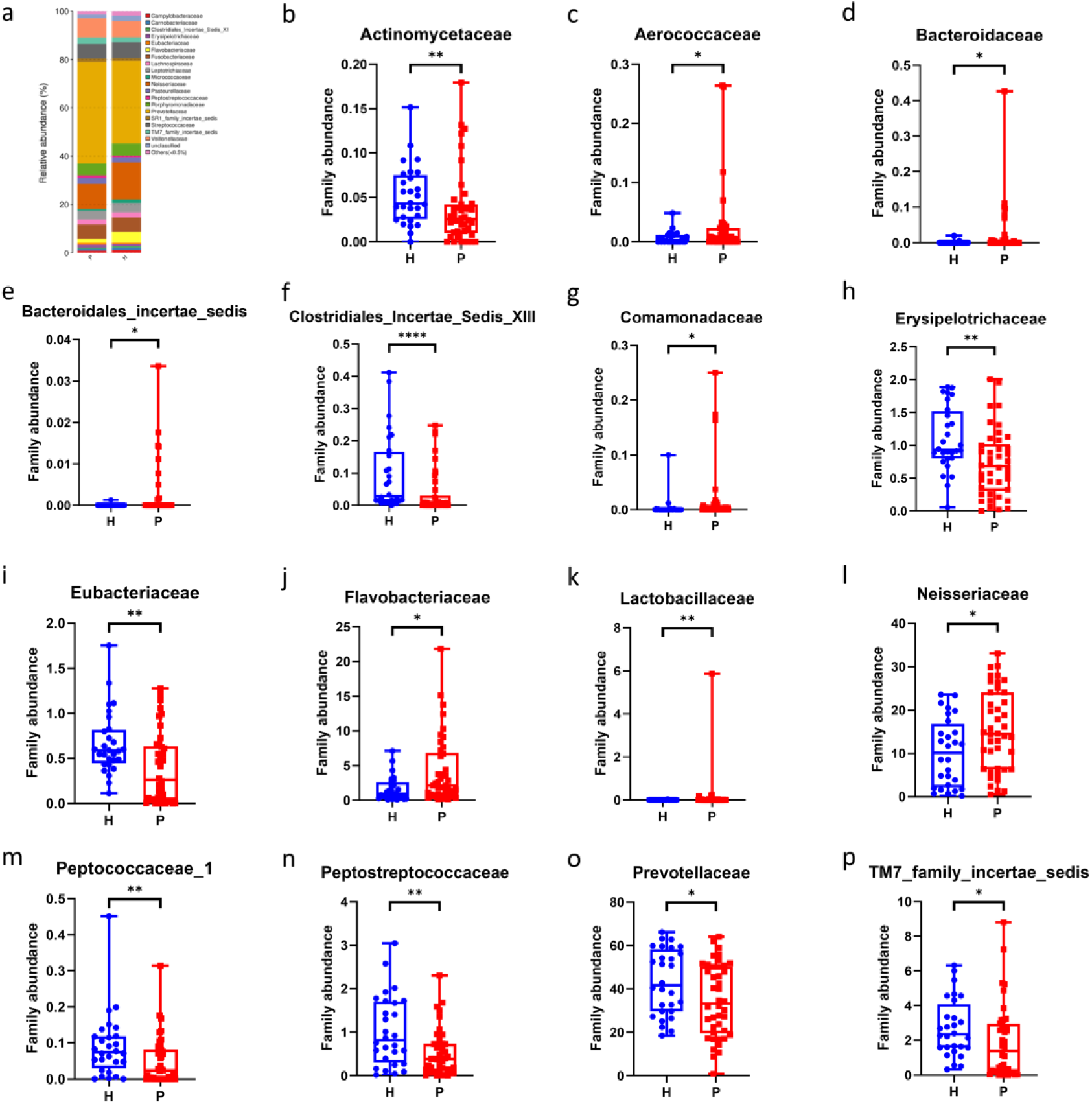
Composition of microbiota at family level and microbes with significantly different abundance. **Note**: \**P* < 0.05; \*\**P* < 0.01; \*\*\**P* < 0.001 and \*\*\*\**P* < 0.0001; Comparison using the Mann–Whitney U test. H= HCs/heathy controls (*n*=28), P= CHF patients (*n*=42).

**Fig.5.**
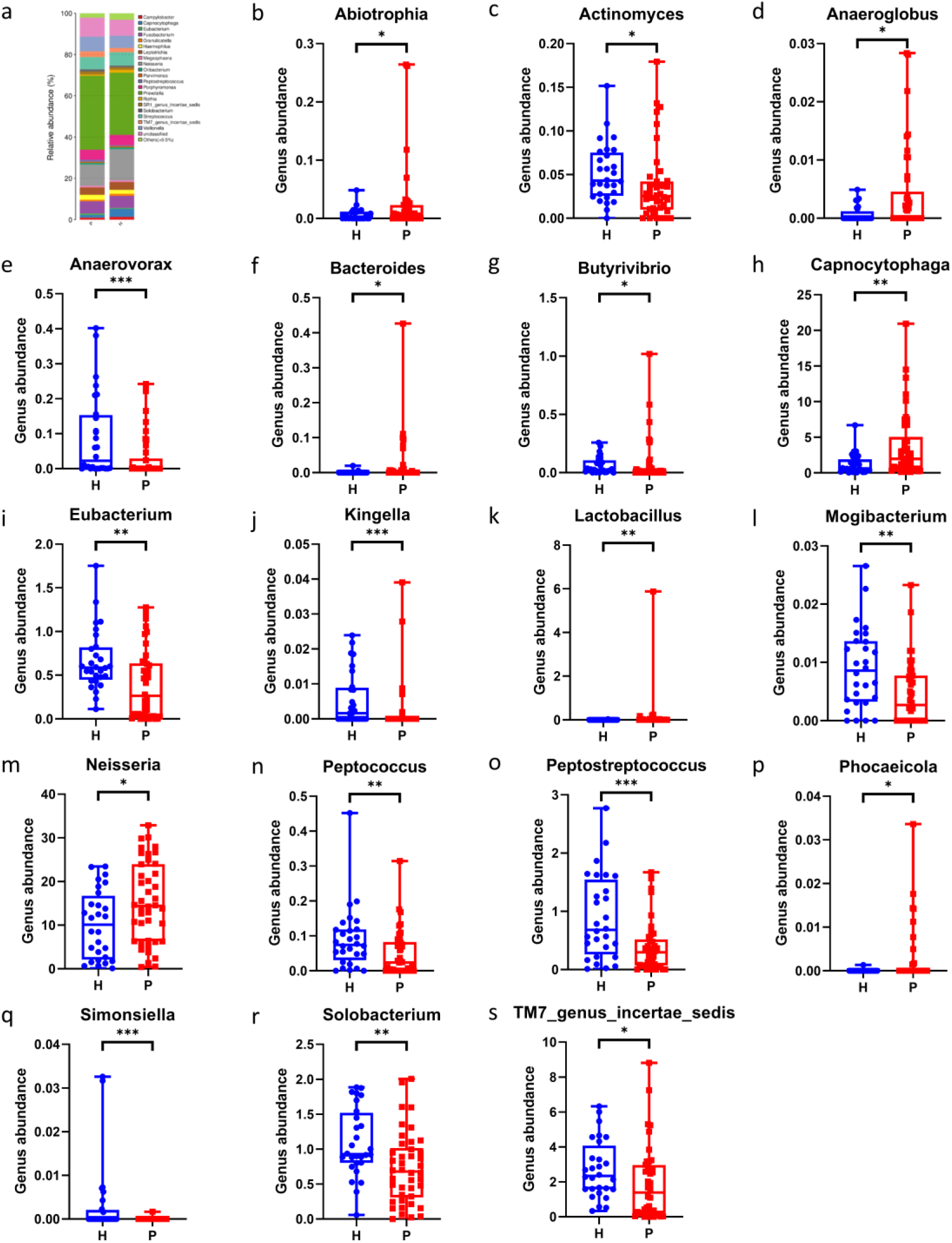
Composition of microbiota at genus level and microbes with significantly different abundance. **Note**: \**P* < 0.05; \*\**P* < 0.01; \*\*\**P* < 0.001 and \*\*\*\**P* < 0.0001; Comparison using the Mann–Whitney U test. H= HCs/heathy controls (*n*=28), P= CHF patients(*n*=42).

**Fig.6.**
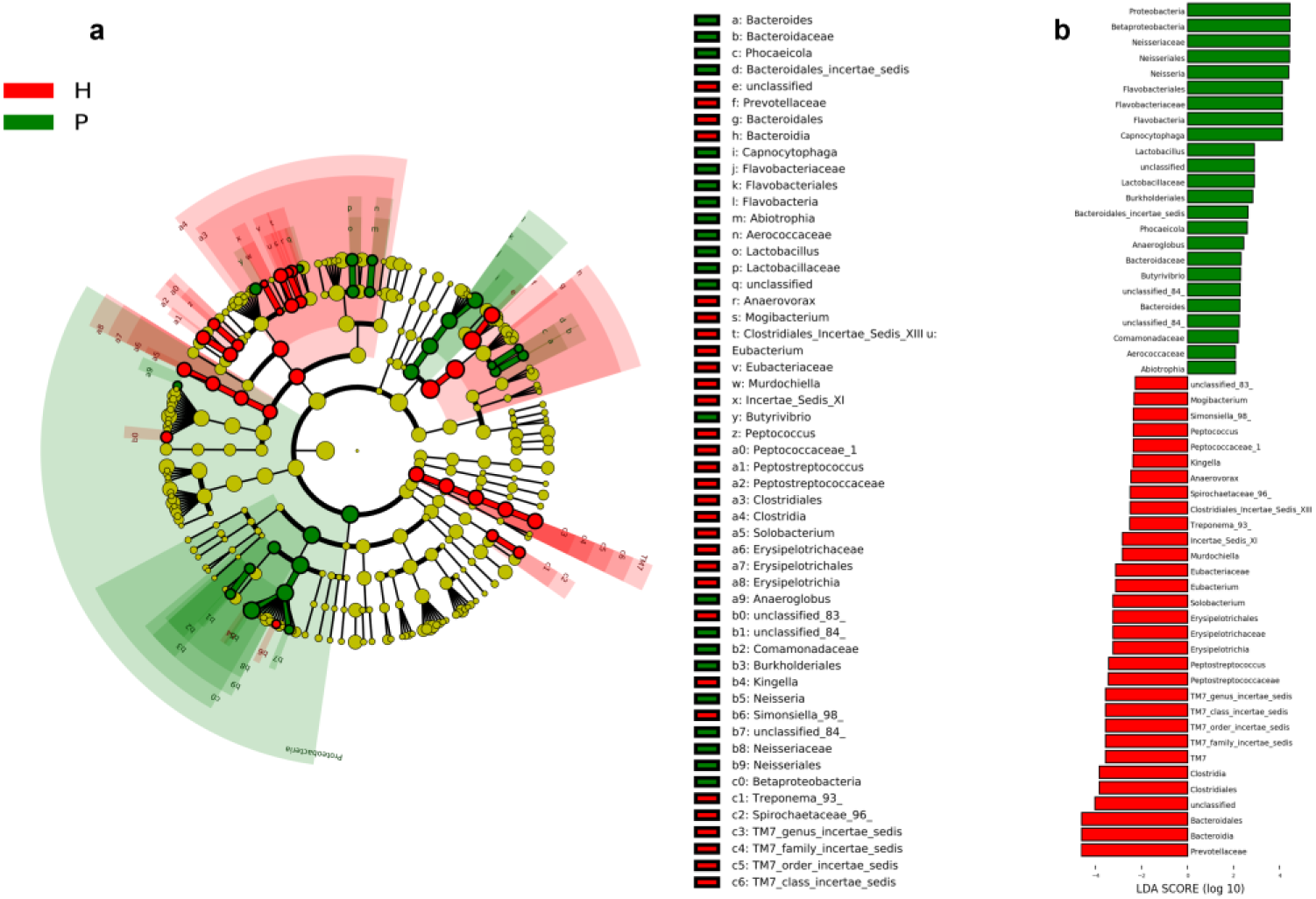
**a**-**b**: LEfSe and LDA analysis based on OTUs characterize microbiota between CHF patients and HCs. **a**: Cladogram using the LEfSe method indicating the phylogenetic distribution of tongue coating microbes associated with patients with CHF (green indicates phylotypes statistically overrepresented in CHF) and healthy controls (red indicates phylotypes overrepresented in healthy controls). Each filled circle represents one phylotype. **b**: Histogram of the linear discriminant analysis (LDA) scores was established for the selected taxa which showed significant difference between the CHF patients and HCs. LDA scores at the log 10 scale are at the bottom. The greater the LDA score is, the more significant the microbial biomarker is in the comparison. **Note**: H= HCs/heathy controls (*n*=28), P= CHF patients (*n*=42).

### 3.4 Diagnostic performance of oral microbiota

Altogether 14 microbes at the genus level could distinguish CHF patients from HCs (Fig.7). *Abiotrophia, Butyrivibrio, Lactobacillus, Capnocytophaga* and *Neisseria* are more abundant in CHF patients, and *Actinomyces, Anaerovorax, Eubacterium, Kingella, Mogibacterium, Peptococcus, Peptostreptococcus, Solobacterium* and *TM7_genus_incertae_sedis* are more abundant in HCs. Furthermore, the AUC of their combined diagnosis was 83.7% (95% confidential interval 74.1%–93.3%)(Fig.7x).

**Fig.7.**
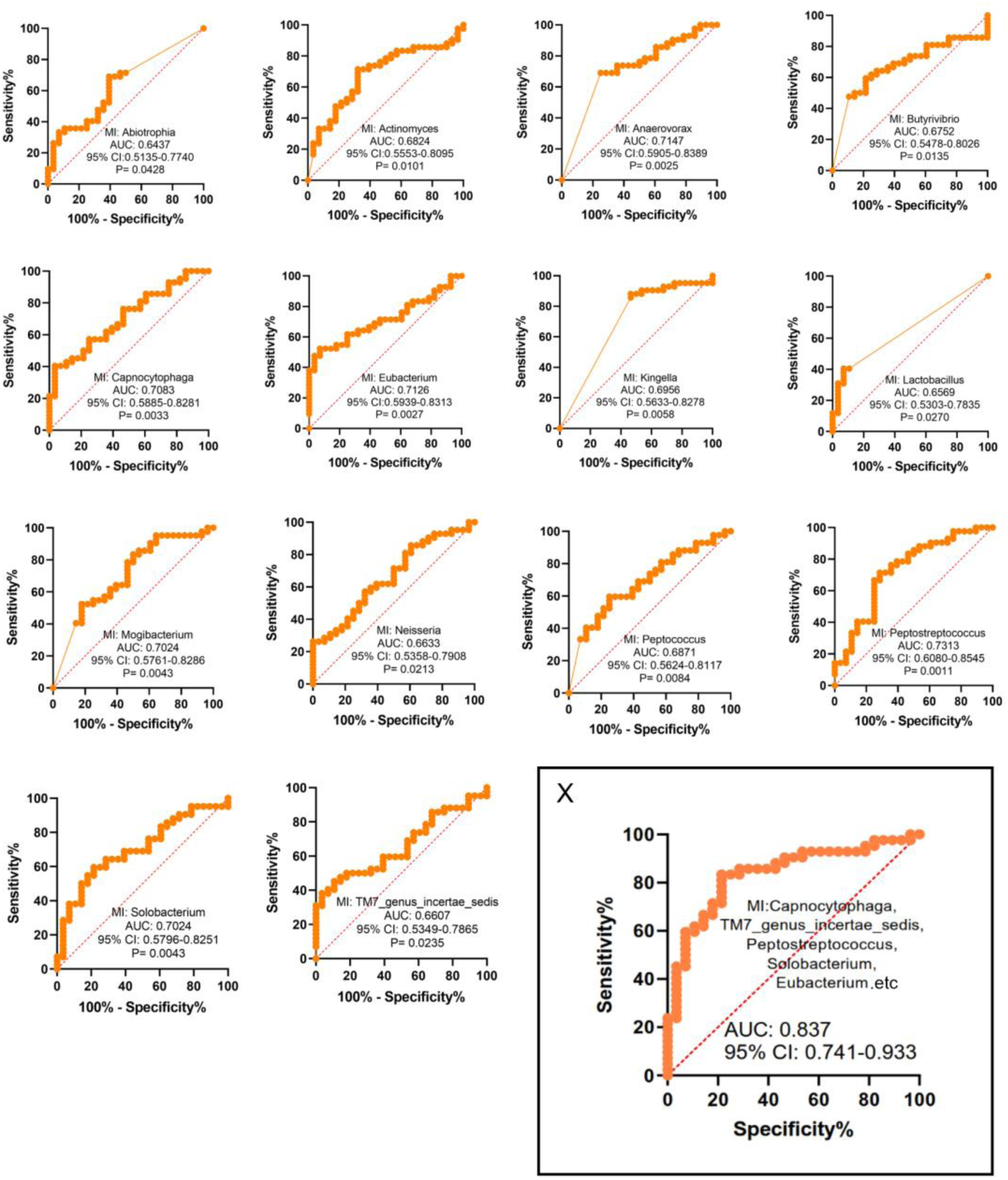
ROC curve of genera with diagnostic significance. **X**: A combination of 14 genera for CHF diagnosis.

#### Identification of genera associated with progression of NYHA Functional Classification

A network was constructed to assess the potential relationship among genera and finally three clusters were obtained (Fig.8(a-c)). The size of the nodes represents the mean abundance of the genera among all samples. *Eubacterium, Solobacterium* and *Rhizobium* were considered as the core genera, and they were compared between groups with different NYHA heart functional grades to evaluate whether they can distinguish the heart functional grade of participants. The abundance of *Eubacterium* and *Solobacterium* in NYHA class II, III and IV were lower than that in HC group. In addition, the abundance of *Eubacterium* and *Solobacterium* seem showed a downward trend with heart functional grade decreasing (Fig.8(e-g)). Nevertheless, although results revealed that oral microbiota links to NYHA class, no significant difference in diversity and quantity of oral microbiota has been found among CHF patients with different classes.

**Fig.8.**
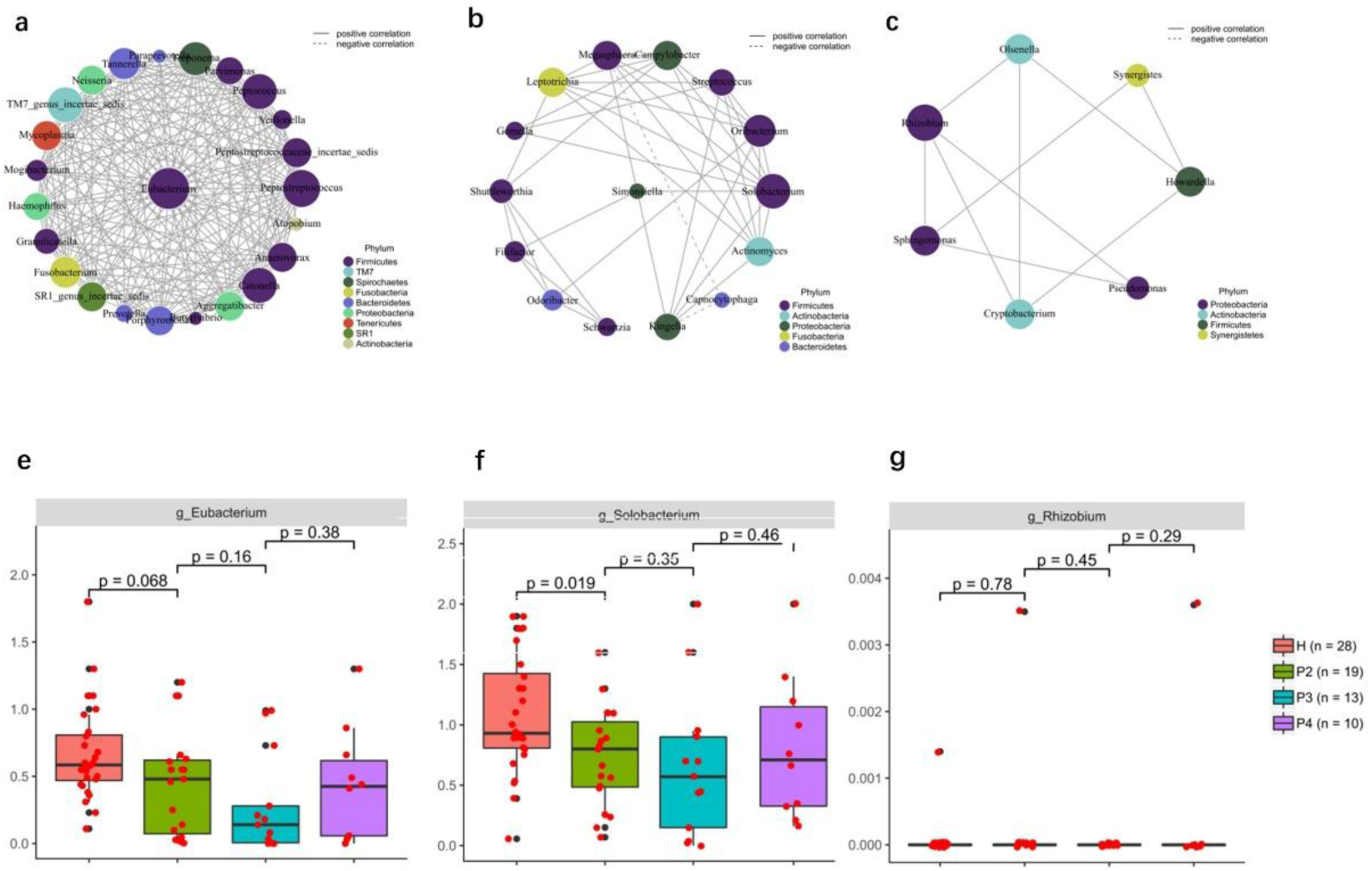
Clusters of core genera of microbes (**a-c**) and the association between three core genera and CHF patients with different heart functional grades (**e-g**). Note: H= HCs/heathy controls(*n*=28), P2= the group of CHF patients with NYHA class II(*n*=19), P3= the group of CHF patients with NYHA class III(*n*=13), P4= the group of CHF patients with NYHA class IV(*n*=10).

## 4. Discussion

In this study, diversity and composition of oral microbiota in CHF patients and HCs was analyzed based on 16S rRNA gene sequencing technology. To the best of our knowledge, it is the first study focused on the association between oral microbiota and CHF. Results showed that oral microbiota of CHF patients was significantly different from that of HCs. Additionally, at different levels, there are various dominant microbes in oral microbiota of CHF patients and HCs respectively.

Chronic heart failure is the final common process as a consequence of various initial cardiac injury and subsequent imbalances in compensatory mechanisms and pathogenic processes[31]. This clinical syndrome involves reduced cardiac output due to impaired cardiac contraction. Typical clinical symptoms of CHF include dyspnoea, fatigue and edema. Lack of validated biomarkers is one of challenges that cannot be ignored in the management of CHF[34]. Though remarkable achievements in treatments have been made over the past few decades, investigation on novel effective biomarkers for the progression and management of CHF is warranted.

There is a prevailing view that oral health serves as an indicator of the state of visceral organs. The coating or moss on the tongue’s surface indicates the current condition of the patient’s metabolism, as well as the presence of toxins and/or morbid, superfluous humors in the body[35], which can reveal etiology and pathogenesis of diseases[36]. Microbiota is capable of ectopic colonization and producing a variety of microbial metabolites which possibly promote cardiovascular events through modulation of pathways associated with energy homeostasis, inflammatory response and immunologic balance[37, 38]. Oral microbiota is mainly comprised of monolayers of sparsely colonized bacteria, bacteria on squamous epithelial cells, structurally complex bacterial entities, and equally free bacteria[39]. The microbiota interacts with tongue papillae, and different species and genera could have same metabolic functions. In general, the diversity of normal oral microbiota is higher than that of patients[40]. There are more consistent reports about oral microbiota at the phylum level than the species level, possibly due to different sampling methods, inclusion criteria, ethnicity, and region[41-43]. A review by Yiwen Li et al suggested the results of 16S rRNA analysis revealed that normal oral colonizing microbiota at the phylum level comprising *Firmicutes, Bacteroidetes, Proteobacteria, Actinobacteria, Spirochaetes, Fusobacteria* and *Synergistetes*[44-46]. Strong evidences existed that oral microbiota is featured with a moderate rate of renewal and good disease sensitivity, making it ideal to investigate [47]. Accordingly, researches focused on oral microbiota are gaining increased attention.

The results of this research revealed 14 types of microbes (Fig. 7), belonging to the lowest taxonomic level in the classification we studied, known as the genus level.They could distinguish CHF patients from HCs and exhibit higher reliability for combined diagnostic significance (Fig. 7X). According to the resluts, *Abiotrophia, Capnocytophaga, Neisseria, Butyrivibrio* and *Lactobacillus* are more abundant in CHF patients (Fig.3g). Previous studies have shown that they are closely related to CHF or other cardiovascular diseases. *Abiotrophia defectiva,* one of species of *Abiotrophia*, is a Gram-positive, non-motile, facultative aerobe having been frequently found in dental plaque[48]. It is a virulent bacterium preferentially affecting endovascular structures and is implicated in many endocarditis cases[49]. Notably, it can cause heart valve destruction resulting in heart failure and excessive embolization rates[50, 51]. *Capnocytophaga* is considered as an opportunistic pathogen. Reports suggested that it can cause various infections including endocarditis, pericardial abscess or coronary heart disease[52-55]. *Neisseria*, a large genus of Gram-negative bacteria[56], colonizes the mucosal surfaces of numerous animal species. Dima Youssef et al suggested *Neisseria elongata* can act as an aggressive organism resulting in serious infections such as infective endocarditis[57]. *Butyrivibrio* is a class of anaerobic bacteria capable of producing butyrate, it can breaks down fiber and sugars through metabolic pathways to produce butyrate. Butyrate has been demonstrated that can exert multiple atheroprotective effects through its anti-inflammatory, anti-proliferative, and lipid-modulating activities[58].*Lactobacillus* is a genus of gram-positive, aerotolerant anaerobes or microaerophilic, non-spore-forming bacteria. *Lactobacillus* exhibits a mutualistic relationship with the human body, as it protects the host against potential invasions by pathogens, and in turn, the host provides a source of nutrients[59]. Although *Butyrivibrio* and *Lactobacillus* has been proved to be beneficial for heart health, their increased abundance in CHF patients may be related to disease compensation, or excessive accumulation could have adverse effects. The underlying mechanisms require further investigation.

On the other hand, *Actinomyces, Anaerovorax, Eubacterium, Kingella, Mogibacterium, Peptococcus, Peptostreptococcus, Solobacterium* and *TM7_genus_incertae_sedis* are more abundant in HCs (Fig.3g). They are among normal oral colonizing microbiota or even can protect against cardiovascular disease including CHF. *Actinomyces* is a genus of *actinomycetes*. Involvement of the heart by actinomycetes is rather rare, and so is endocardial, valvular, and myocardial involvement which is usually secondary to pericardial disease[60-62]. *Kingella* are Gram-negative organisms that colonize the human respiratory tract. Although we found *Kingella* were of more abundance in HCs in this study, they can still lead to some diseases, which causes endocarditis and bacteremia[63]. Among *Kingella* species, *Kingella kingae* is the most frequent human pathogen[64]. *Kingella* endocarditis has been reported in all age groups, especially in children, which may involve native or prosthetic valves[65-67]. *Mogibacterium* is a Gram-positive, strictly anaerobic and non-spore-forming bacterial genus from the family of *Eubacteriaceae* [68]. *Peptococcus* is a genus of Gram-positive, anaerobic, coccoid bacteria; they are part of the flora of the mouth, upper respiratory tract and large intestine[69]. Jiyeon Si et al reproted that validation of identified oral microbiota by quantitative PCR (qPCR) revealed that healthy controls possessed a higher abundance of *Peptococcus* compared with metabolic syndrome group[70].*TM7_genus_incertae_sedis* is a bacteria from the *Saccharibacteria* phylum, whic are ubiquitous members of the human oral microbiome, and accumulating evidence links their association with periodontal disease[71].

It needs to be stressed we found the abundance of *Eubacterium* and *Solobacterium* in CHF patients were significantly lower than those of HCs and displayed downward trends as NYHA class increases (Fig.8), which may be useful for distinguishing cardiac function grades by NYHA class in patients with CHF in the clinic. *Solobacterium*, a Gram-positive, strictly anaerobic, and non-spore-forming bacterium[72], has been identified as a ubiquitous component of the human microbiota, colonizing multiple anatomical sites[73]. *Eubacterium*, also known as bacteria, is a genus of Gram-positive bacteria in the family *Eubacteriaceae*. Members of the genus *Eubacterium*, a part of the core human gut microbiota, are among a new generation of potentially beneficial microbes. *Eubacterium spp.* carry out bile acid and cholesterol transformations in the gut, thus contributing to their homeostasis[74]. It is reported that butyrate producers in the gut of humans and mice including *Eubacterium spp.* show a significant negative correlation with atherosclerotic cardiovascular disease[75-77]. Gut metagenomes analysis from patients with atherosclerosis displays a reduction of butyrate producers including *Eubacterium spp.* and *Roseburia spp.* in contrast with HCs[75, 77]. This protective mechanism can be attributed to the anti-atherogenic properties of butyrate. Chan YK et. al. revealed that *Eubacterium spp.* and other butyrate producers were negatively associated with plasma cholesterol, MMP-9 and A-FABP (biomarkers for cardiovascular pathologies) in mice with atherosclerosis[76]. Moreover, *Eubacterium spp.* can transform cholesterol in human body and have hypocholesterolemic effect, thereby they are capable of providing protection against cardiovascular diseases. Collectively, as one of butyrate producers, *Eubacterium spp.* may be potential to restore a dysbiotic gut microbiota and modulate inflammation in subjects with acute cardiovascular disease and shed light on the diagnosis and treatment of cardiovascular diseases including CHF.

In addition, it is worth exploring that we found the abundance of *Proteobacteria* (also known as *Pbac*) were significantly higher in CHF patients (P<0.05) in this study. According to a previous study by Amar et al., blood microbiota dysbiosis and especially increase of *Proteobacteria* were associated with the onset of cardiovascular events in a general population after adequate adjusting for known cardiovascular risk factors, such as smoking, thus *Proteobacteria* was shown to be independent markers of the risk of cardiovascular disease[78]. Nevertheless, Kamo T et al implied that significant differences were not observed between the samples from HCs and heart failure patients in terms of relative abundances of *Proteobacteria*.[79] In addition, older heart failure patients had larger quantities of *Proteobacteria* compared with younger ones[79]. Based on it, we hold the opinion that the difference of the abundances of *Proteobacteria* in CHF patients and HCs requires further investigation, and it cannot yet be used in the differentiation of CHF.

In summary, the current evidence demonstrates a bidirectional relationship between heart failure and oral microbiota. Heart failure significantly impacts the abundance and diversity of microbial communities in the oral cavity, and dysbiosis of the oral microbiota serves as a robust biomarker for the progression of heart failure. Specifically, conditional pathogens may contribute to heart failure pathogenesis through multiple mechanisms, including systemic infections, pro-inflammatory responses, valve destruction and the presence of multiple emboli. Conversely, bacterial species that are more abundant in healthy individuals appear to exert cardioprotective effects through various pathways, such as anti-inflammatory actions, ventricular remodeling modulation, metabolic disorder amelioration, and lipid profile regulation. It is also noteworthy that we propose that the pathological process of CHF has an impact on oral microbiota, making normal microbiota such as *Eubacterium* and *Solobacterium* display irregular features in patients with different NYHA classes as a result of microbiota dysbiosis.

Furthermore, these microbial communities may play a pivotal role in the pathophysiological interplay between chronic heart failure (CHF) and its frequent comorbidities, including type 2 diabetes mellitus and glomerulonephritis. This microbial-mediated crosstalk potentially offers novel therapeutic targets and diagnostic approaches for managing these complex chronic disease interactions, thereby providing new perspectives for the development of integrated treatment strategies in multimorbidity management.

Nevertheless, there are several limitations about this research. Firstly, given that the majority of CHF patients are elderly, a considerable age disparity exists between the HCs and CHF patients included in this study. Age-related variations in the composition of oral microbiota may influence the findings, potentially introducing an age-dependent bias into the results. Secondly, we just made a comparison about features of oral microbiota between CHF patients and HCs, the underlying molecular biological mechanisms concerning oral microbiota and CHF still requires further investigation. Thirdly, the study sample is limited to two hospitals, and the sample size is relatively small, so generalizing the findings to a larger population group should be done with caution. Given that the composition and activity of oral microbiota depends on many factors, such as dietary, environment, ethnic groups and so on, findings in this single-center study need to be validated in a variety of population with lager sample sizes. Fourth, this research was conducted without microbial metabolomics analysis. Microbial metabolomics detection techniques can clarify the alteration of oral metabolites, which may be helpful to reveal the microecosystem network of oral microbiota dysbiosis.

## Conclusion

CHF is the final stage in several heart diseases with increasing prevalence and incidence. Currently, the diagnosis of CHF still remains a challenge. Through the analysis of oral microbiota in CHF patients and HCs, this study suggested the dysbiosis of the oral microbiota in CHF patients and revealed potential biomarkers for CHF diagnosis. Altogether 14 microbes could distinguish CHF patients from HCs. *Eubacterium, Solobacterium* and *Rhizobium* were three core genera and when the abundance of *Eubacterium* and *Solobacterium* exhibited downward trends NYHA class increases. Collectively, oral microbiota is potential to be biomarkers associated with CHF, which could be objective, non-invasive and suitable for long-term monitoring. This study provides a more solid foundation for advancing clinical diagnostic practice and experimental evidence for CHF management. This groundbreaking research significantly contributes to the elucidation of chronic heart failure (CHF) pathophysiology while establishing a robust foundation for the development of novel therapeutic strategies. Furthermore, emerging microbiota-based diagnostic modalities demonstrate substantial potential to revolutionize current clinical paradigms in CHF management.

## Abbreviations

CHF: chronic heart failure
HCs: healthy controls
NYHA: New York Heart Association
NT-proBNP: N-terminal pro-b-type natriuretic peptide
LEfSe: Linear discriminant analysis effect size
OTUs: operational taxonomic units
AUC: Area Under the Curve
TMAO: trimethylamine N-oxide.

## Acknowledgments

The authors wish to thank all the participants in this study.

## Authors’ Contributions

Xing Ye: Methodology, Data curation, Formal analysis, Visualization, Original Draft; Jie Ja: Software, Visualization, Investigation, Original Draft; Xiaotao Jiang: Software, Visualization, Investigation, Original Draft; Jiao Jiao Li:Resources, Investigation; Yanlin Zeng: Resources, Investigation; Yan Wang: Resources, Investigation; Zhongqi Yang: Review & Editing, Supervision; Jing Sun: Review & Editing, Supervision; Tianhui Yuan:Conceptualization, Project administration, Review & Editing. All authors read and approved the final manuscript.

## Declarations

### Ethics approval and consent to participate

This study was approved by the Institutional Review Board of The First Affiliated Hospital of Guangzhou University of Chinese Medicine (No.K[2020]061). The study was performed according to the Helsinki Declaration and Rules of Good Clinical Practice. All participants approved and signed informed consents upon enrollment. Please refer to the Supplementary materials for ethical approval.

### Consent for publication

Not applicable.

### Author details

*^1^The First Clinical Medical College, Guangzhou University of Chinese Medicine, Guangzhou, China; ^2^ The First Affiliated Hospital of Guangzhou University of Chinese Medicine, Guangzhou, Guangdong, China; ^3^ Kolling Institute, Faculty of Medicine and Health, The University of Sydney, St Leonards, NSW 2065, Australia; ^4^ School of Biomedical Engineering, Faculty of Engineering and IT, University of Technology Sydney, Sydney, NSW 2007, Australia; ^5^ School of Acupuncture and Rehabilitation, Guangzhou University of Chinese Medicine, Guangzhou, China; ^6^ Department of Geriatrics, The First Affiliated Hospital of Guangzhou University of Chinese Medicine, Guangzhou, China; ^7^ Department of Cardiology, International Medical Services, The First Affiliated Hospital of Guangzhou University of Chinese Medicine, Guangzhou, China; ^8^ Rural Health Research Institute, Charles Sturt University, New South Wales, Australia; ^9^ Data Science Institute, University of Technology Sydney, Sydney, NSW 2007, Australia; ^10^ School of Health Sciences and Social Work, Griffith University, QLD 4215, Gold Coast*

